# Assessing HD-EEG functional connectivity states using a human brain computational model

**DOI:** 10.1101/2021.10.14.464481

**Authors:** Judie Tabbal, Aya Kabbara, Maxime Yochum, Mohamad Khalil, Mahmoud Hassan, Pascal Benquet

## Abstract

**Objective:** Electro/Magnetoencephalography (EEG/MEG) source-space network analysis is increasingly recognized as a powerful tool for tracking fast electrophysiological brain dynamics. However, an objective and quantitative evaluation of pipeline steps is challenging due to the lack of realistic ‘controlled’ data. Here, our aim is two-folded: 1) provide a quantitative assessment of the advantages and limitations of the analyzed techniques and 2) introduce (and share) a complete framework that can be used to optimize the entire pipeline of EEG/MEG source connectivity.

**Approach:** We used a human brain computational model containing both physiologically based cellular GABAergic and Glutamatergic circuits coupled through Diffusion Tensor Imaging, to generate high-density EEG recordings. We designed a scenario of successive gamma-band oscillations in distinct cortical areas to emulate a virtual picture-naming task. We identified fast time-varying network states and quantified the performance of the key steps involved in the pipeline: 1) inverse models to reconstruct cortical-level sources, 2) functional connectivity measures to compute statistical interdependency between regional signals, and 3) dimensionality reduction methods to derive dominant brain network states (BNS).

**Main Results:** Using a systematic evaluation of the different decomposition techniques, results show significant variability among tested algorithms in terms of spatial and temporal accuracy. We outlined the spatial precision, the temporal sensitivity, and the global accuracy of the extracted BNS relative to each method. Our findings suggest a good performance of wMNE/PLV combination to elucidate the appropriate functional networks and ICA techniques to derive relevant dynamic brain network states.

**Significance:** We suggest using such brain models to go further in the evaluation of the different steps and parameters involved in the EEG/MEG source-space network analysis. This can reduce the empirical selection of inverse model, connectivity measure, and dimensionality reduction method as some of the methods can have a considerable impact on the results and interpretation.

## Introduction

The human brain is currently recognized as a complex network in which spatially separated brain regions are functionally and dynamically communicating during tasks (Bola and Sabel, 2015; Hassan et al., 2015; O’Neill et al., 2017) and at rest (Allen et al., 2014; Baker et al., 2014; Kabbara et al., 2017). The characterization of transient (dynamic) networks is of utmost importance to better understand the brain processes in healthy brains, as well as in neurological disorders (Kim et al., 2017; Sitnikova et al., 2018). Hence, the analysis of large-scale dynamic functional connectivity (dFC) has become a key goal in cognitive and clinical neuroscience. Among existing neuroimaging modalities, electro/magneto-encephalography (EEG/MEG) has major benefits in exploiting the spatiotemporal organization of the brain, since it provides a non-invasive measure of electrical activity at the -sub-millisecond timescale (Brookes et al., 2018; Vidaurre et al., 2018).

The ‘EEG/MEG source connectivity’ is a potential tool to identify brain networks with high space/time resolution at the cortical space through sensor-level signals (de Pasquale et al., 2010; Hassan et al., 2014; Hassan and Wendling, 2018; Schoffelen and Gross, 2009). This technique is mainly based on two steps: (1) source reconstruction and (2) functional connectivity. An additional (3) third step is crucial to apply to group the set of hundreds (or even thousands) of functional brain networks that fluctuate over the whole time recording into dominant ‘Brain Network States (BNS)’ that describe essential brain patterns activities (O’Neill et al., 2018; Tait and Zhang, 2021). Yet, there is no agreement on the best (if any) source reconstruction/connectivity/clustering algorithms to use when studying EEG/MEG dynamic networks.

This is partially due to the fact that a quantitative evaluation of the various methods’ robustness is still missing, as ‘ground truth’ is almost impossible to get when studying empirical data. Therefore, validating the above-mentioned ‘three-steps’ pipeline using simulated EEG data is needed. In this context, some studies used a toy model where brain sources activity is modeled by oscillatory sinusoidal and Gaussian functions (Halder et al., 2019) or multivariate autoregressive (MVAR) filters with volume conductor head models generating pseudo-EEG data (Anzolin et al., 2019; Haufe and Ewald, 2019). However, such models are too simplistic compared to the complexity of real brain activity. On the other hand, few studies executed a preliminary performance quantification of dimensionality reduction using a set of pre-defined connectivity matrices, considered as simulated brain networks without any constraint on how these matrices are computed (Kabbara et al., 2019; O’Neill et al., 2017; Tabbal et al., 2021; Zhu et al., 2020). Here, we address these challenges by simulating the whole brain EEG dynamics by building physiologically inspired networks based on a realistic neural mass model using a human brain computational model, named ‘COALIA’ (Bensaid et al., 2019). It contains several subtypes of GABAergic neurons (VIP- SST- and PV-neurons) and Glutamatergic (Pyramidal cells) neurons coupled through Diffusion Tensor Imaging (DTI) based structural connectivity, to simulate realistic HD-EEG data

Technically, EEG simulated signals were computed via the forward model as previously shown by recent works (Allouch et al., 2020; Bensaid et al., 2019). In this paper, we aim to investigate to what extent we can effectively track time-varying brain networks that dominate neuronal activity using HD-EEG scalp signals, specifically on a short timescale that is commensurate with cognitive tasks. To this end, we benefit from the COALIA model to simulate a set of dynamic brain network states (BNS) related to a fast timescale cognitive task (picture naming). For this purpose, coherent gamma oscillations were produced in cortical areas of the ventral visual pathway following a picture naming dynamic scenario, as observed in real human data (Hassan et al., 2015).

The Local Field Potentials (LFP) of 20 subjects were simulated and the corresponding HD-EEG signals were calculated. In this context, we evaluated the performance of some frequently used inverse/connectivity algorithms (weighted Minimum Norm Estimate (wMNE), Exact low-resolution brain electromagnetic tomography (eLORETA) / Phase Locking Value (PLV), weighted Phase Lag Index (wPLI), Amplitude Envelope Correlation (AEC)). In addition, we quantified the efficiency of many common source separation methods including three variants of Independent Component Analysis (ICA), Principal Component Analysis (PCA), Non-negative Matrix Factorization (NMF), and Kmeans. To our knowledge, no previous study has validated the performance of this ‘three-steps’ pipeline with the existence of ground truth provided by realistic HD-EEG simulation. We hope that this paper can be used as a benchmark for researchers who intend to investigate methodological or experimental issues to optimize EEG/MEG dynamic networks analysis.

## Methods

The full pipeline of our study is divided into six steps as summarized in Figure 1.

**Figure 1.**
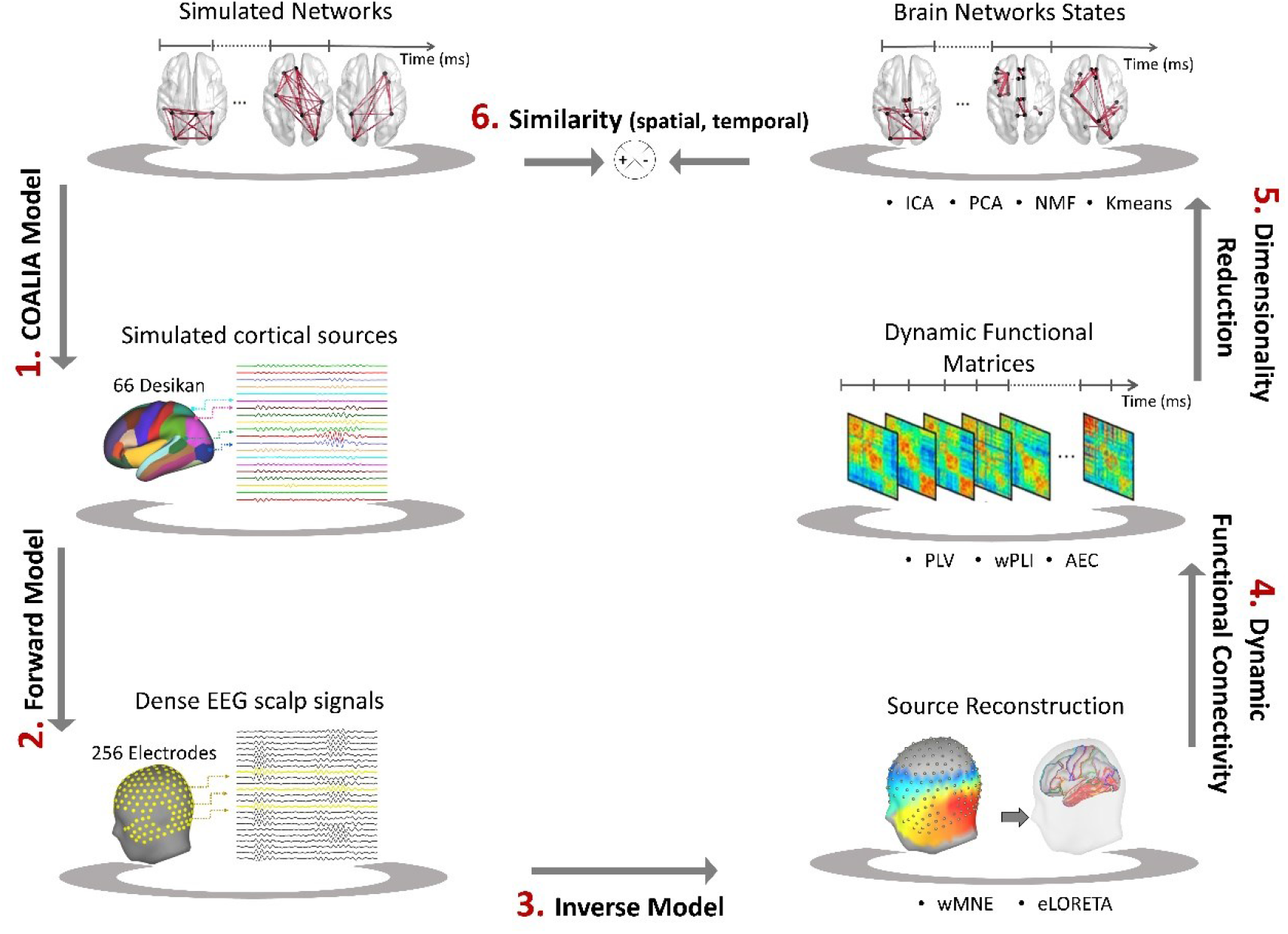
Structure of the investigation. (1) COALIA Model was used to simulate cortical sources from a set of simulated networks. (2) 257 scalp level signals were generated using the forward model. (3) Cortical-level sources (66 Desikan regions) were reconstructed using wMNE and eLORETA methods. (4) Dynamic Functional Connectivity between sources was calculated using PLV, wPLI, and AEC methods. (5) Dimensionality reduction methods including ICA, PCA, NMF, and Kmeans were applied to reconstruct dominant brain network states. (6) Spatial and Temporal similarities were finally calculated to measure the performance of brain network estimation.

### 1. The ‘upgraded’ COALIA Model

#### 1.1. Simulations

The simulated cortical-level activity was generated using an updated version of the computational model named COALIA. It is based on the interconnected Neural Mass Model (NMM) respecting human structural connectivity based on DTI. This model has proven its ability to produce realistic EEG when compared to real EEG recordings obtained in humans during awake and deep sleep (Bensaid et al., 2019). In their work, Bensaid et al. obtained similar morphology, spectral content, and topographical voltage distribution between simulated and real EEG data.

Briefly, each microcircuit based on neural mass can produce distinct brain oscillations such as alpha-rhythms through the Pyramidal somatostatin/positive (SST) loop, gamma-rhythms through Pyr-parvalbumin (PV) loop, delta-rhythms through increased thalamocortical connectivity and disinhibition through VIP-SST microcircuits. Therefore, using distinct local oscillations cortical-level simulated signals taking into account macro and micro-circuitry of the human cortex, it becomes possible to generate coherent oscillations in different brain regions of interest to simulate dynamic functional connectivity during a cognitive task.

i. At the local level, each neural mass consists of subsets of a glutamatergic pyramidal neuron population and three types of GABAergic interneuron populations (VIP-, PV-, SST-interneurons) with physiologically based kinetics (fast/slow) and interconnection between neural models. Increased excitability of the perisomatic targeting PV-interneurons onto pyramidal neurons produced gamma oscillations (30-45Hz). Increased excitability of the dendritic targeting SST-interneurons onto pyramidal neurons produced alpha oscillations (8-12Hz). Remote activation of the VIP-interneurons onto SST induced disinhibition and strengthened coherent gamma oscillations between coactivated interconnected cortical areas.
ii. Then, at the global level, the large-scale model is constructed based on *nROIs*=66 regions of interest from the standard anatomical parcellation of the Desikan-Killiany atlas (Desikan et al., 2006) (right and left insula were excluded, leaving 66 brain regions). In this case, each neural mass represents the local field potential (LFP) of one atlas region, in which the activity is assumed to be homogeneous. The template brain morphology (Colin) is used to spatially distribute neural masses over the cortex. Neural masses are synaptically connected through long-range glutamatergic projections among pyramidal neurons and GABAergic interneurons (Bensaid et al., 2019).
iii. In order to improve the realism of simulated functional connectivity, COALIA was upgraded using the structural connectivity matrices averaged among a large number of healthy participants (487 adult subjects) through DTI as provided in the Human Connectome Project (HCP) (https://www.humanconnectome.org/) (Van Essen et al.,

2013). Hence, we defined our large-scale structural connectivity matrix that represents the density of fibers between all pairs of 66 cortical regions. We used the averaged fractional anisotropy measures to connect all neural masses. The time delay was organized in the form of a matrix where the elements represent the Cartesian distance between cortical NMMs divided by the mean velocity of traveling for action potentials. Here, we used a mean velocity of 7.5 m/s.

A schematic overview of the upgraded COALIA model is presented in the Supplementary Materials (Figure S1).

It is worthy to note that, in this study, we aim to use a previously validated model (COALIA) for testing methods included in the functional connectivity states estimation, rather than to introduce or validate a new model. Therefore, the reader can refer to (Bensaid et al., 2019) for a more detailed description of the COALIA model.

#### 1.2. Scenario

As we are interested in tracking task-related dynamic brain network states (BNS), we consider the picture-naming task dynamic scenario, inspired by a previous publication (Hassan et al., 2015). Thus, the input of the COALIA model is a set of six simulated networks consecutive in time from 0 to 535ms. At each time interval T_i_ (i=[1;6]), different regions of interest (ROIs) are activated as detailed in Figure 2. To be active or inactive, the parameters of each neural mass are tuned accordingly to generate background or gamma activity, based on the Desikan-Killiany atlas (Desikan et al., 2006).

**Figure 2.**
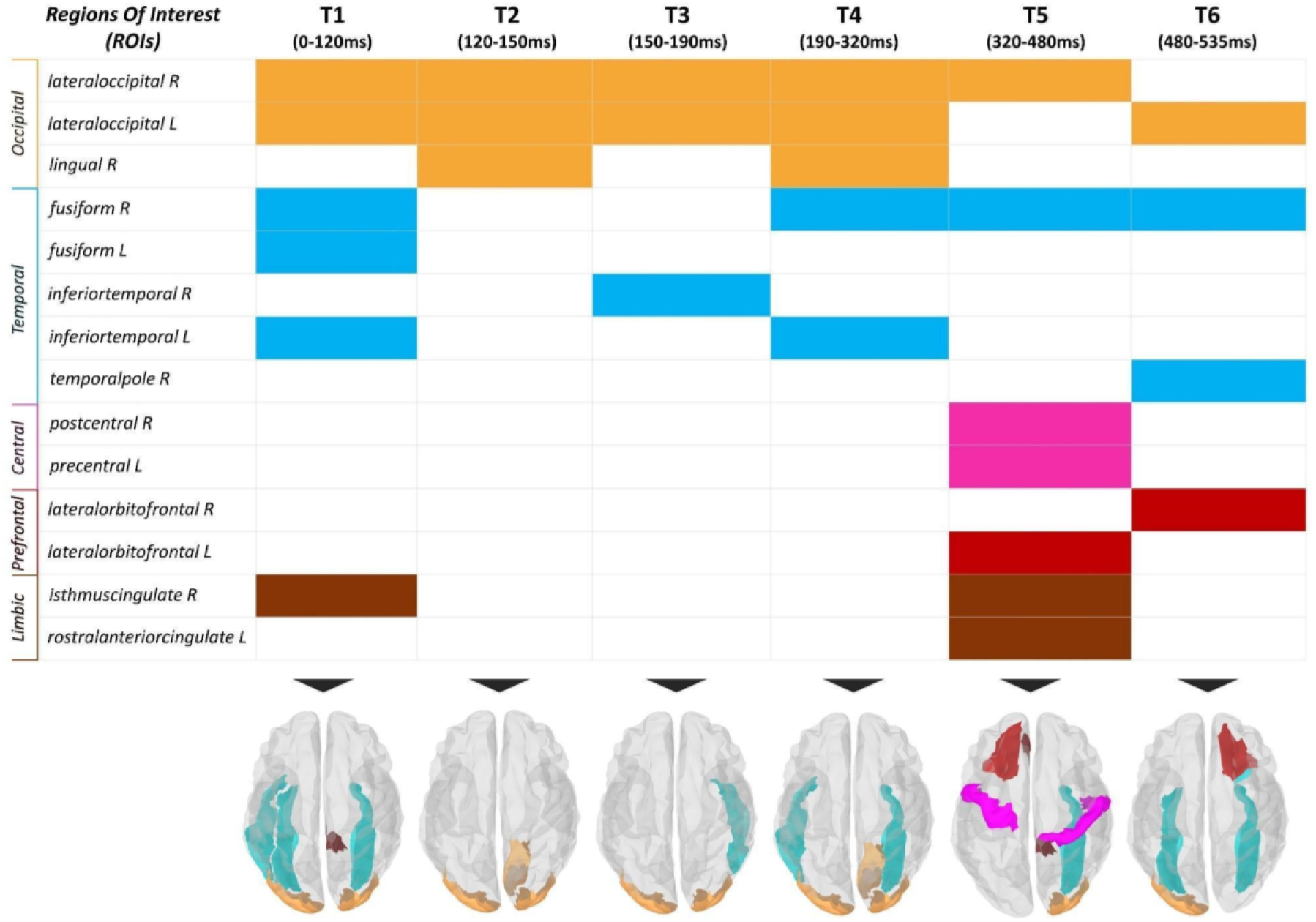
Simulation Scenario. All activated ROIs are shown for each time interval (from T1 to T6). Different color codes refer to different lobe affiliations. Orange color stands for occipital lobe, blue for temporal lobe, pink for central lobe, red for prefrontal lobe, and brown for limbic lobe.

The activity of brain sources is simulated in the gamma band [30-40Hz] as it is shown to be the most relevant frequency band in the context of such cognitive task (Fell and Axmacher, 2011; Rodriguez et al., 1999), with a sampling frequency of f_s_=1024Hz. To perform a group-level brain network analysis, LFP signals are simulated for *nSubs*=20 subjects with *nTrials*=100 trials for each subject. To consider an intra-subjects variability on the EEGs (between the *nTrials)*, the input noise parameters (mean and variance) for all neural masses are randomly adjusted (uniform law) by ± 20% on each simulated LFP. To consider an inter-subject variability (between the *nSubs)*, each value of the connectivity matrix, used to link the 66 neural masses, has been randomly (uniform law) increased or decreased by 10%. Each LFP trial lasts for 2 seconds; including 1-second pre-stimulus and 1-second post-stimulus.

We checked that every neural mass involved in the scenario (Figure 2) generates gamma oscillations and that co-activated neural masses synchronized as measured by the increased PLV during the periods defined by the scenario. Simulated data are available on GitHub (https://github.com/judytabbal/dynCOALIA).

### 2. Forward Model

In order to generate simulated EEG data (‘*X_t_*’) from simulated cortical time series (‘*S_t_*’), we solved the forward problem as follows:

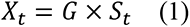

where ‘*G*’ is the lead field matrix to be computed, which describes the electrical and geometrical characteristics of the head. To this end, T1 magnetic resonance imaging MRI (Colin27 template brain, (Holmes et al., 1998)) was spatially aligned with head coordinates and segmented. Thus, realistically shaped shells representing brain, skull, and scalp surfaces were prepared to build a realistic volume conduction head model using the Boundary Element Method (BEM) provided by the OpenMEEG package (Fieldtrip, (Gramfort et al., 2010)). In this work, we used 257 electrodes density (EGI, Electrical Geodesic Inc.) for EEG electrode configuration. Therefore, the computed lead field matrix ‘*G*’ (dimension: 66 × 257) describes the physical current propagation from the sources located at centroids of the 66 brain atlas regions to the 257 EEG sensors. The corresponding source orientation was constrained to be normal to the surface.

Finally, a spatially and temporally uncorrelated white noise was added to the scalp EEG signals to mimic measurement noise (Anzolin et al., 2019).

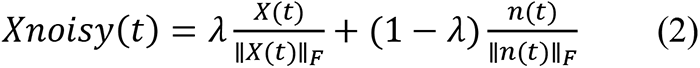

Where *X*(*t*) are the scalp EEG signals and *n*(*t*) is the white uncorrelated noise. ‖‖*_F_* correspond to the Frobenius norm and λ determine noise level added (signal to noise ratio: SNR) varied between 0.7 and 1 (no added noise) with 0.05 step.

### 3. Inverse Solutions

Next, the temporal dynamics of the cortical sources were reconstructed by solving the inverse problem. It consists of evaluating the parameters of the template source space, including position, orientation, and magnitude of current dipoles. As in the forward model, we constrained the position of cortical sources at the centroid locations of 66 Desikan-Killiany regions, and the orientation to be normal to the cortical surface. Therefore, dipole moment at time t (‘*S_t_*’) can be calculated from sensor EEG time series ‘*X_t_*’ as follows:

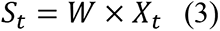

Where ‘W’ denotes the inverse matrix, often called spatial filters (or weights). Algorithms to estimate ‘W’ can be divided into beamforming and least-squares minimum-norm type estimates. In this study, we are interested in evaluating inverse methods based on the latter type as they are widely used in EEG source connectivity analysis. In particular, we selected two methods: (1) weighted minimum norm estimate (wMNE) and (2) exact low-resolution electromagnetic tomography (eLORETA). These two methods mainly differ in the prior assumptions of the source covariance as explained below.

#### 3.1. Weighted Minimum Norm Estimate (wMNE)

The weighted minimum norm estimate (wMNE) (Hämäläinen and Ilmoniemi, 1994) searches for a solution that fits measurements with a least square error. It compensates for the bias of the classical minimum norm estimate (MNE) of favoring weak and surface sources. Technically, depth weighting is implemented by introducing a diagonal weighting matrix *B* in order to boost the impact of deep sources:

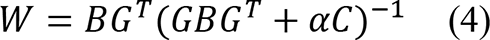

Where ‘*G*’ is the lead field matrix, ‘α’ is the regularization parameter (inversely proportional to signal to noise ratio and set to 0.3 in our case) and ‘*C*’ is the noise covariance matrix (computed from one-second pre-stimulus baseline). ‘*B*’ is the source covariance matrix used to adjust the properties of the solution by incorporating some prior knowledge about the spatial distribution of the source activity. ‘*B*’ is inversely proportional to the norm of lead field vectors.

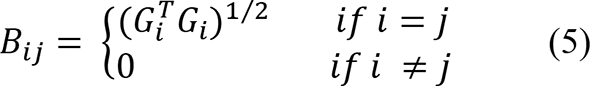

In this work, we used the Matlab function implemented in Brainstorm toolbox codes (https://neuroimage.usc.edu/brainstorm) to compute wMNE (Tadel et al., 2011), with the signal to noise ratio set to 3 and depth weighting value to 0.5 (default values).

#### 3.2. Exact low-resolution brain electromagnetic tomography (eLORETA)

The exact low-resolution electromagnetic tomography (eLORETA) (Pascual-Marqui, 2007) is a genuine inverse solution that gives more importance to the deeper sources with reduced localization error despite the presence of measurement and structured biological noise.

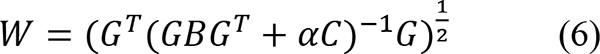

Here, eLORETA was evaluated using the FieldTrip toolbox (http://www.fieldtriptoolbox.org) following the approach of (Pascual-Marqui, 2007) with the default regularization parameter set to 0.05.

#### 3.3. Source Space Resolution

When solving the forward/inverse problems, various techniques can be applied based on different source-space resolutions. One approach is to calculate the lead field vectors within a cortical mesh of high resolution (15000 vertices in most cases), followed by a projection onto an anatomical framework where all lead field vectors belonging to a common atlas region are averaged (Hassan et al., 2014). In this case, the inverse problem is ill-posed (15000 cortical sources>>257 EEG sensors). Another approach is to directly select the centroids of 66 Desikan regions where the lead field vectors are calculated. In our simulation study, we addressed this technical point and tested both approaches to assess the impact of source-space resolution used.

### 4. Dynamic Functional Connectivity (dFC)

We estimated the functional connectivity between reconstructed regional time series. As we aim to undertake the dynamics of brain states, we used a sliding window approach (*window.size = δ, window.step = Δ*), where connectivity is measured within each temporal window. In the case of phase synchronization and based on (Lachaux et al., 2000), the smallest number of cycles recommended to have a compromise between good temporal resolution and good accuracy is 6. Thus, as we are working in the gamma band (*central frequency* =35*Hz*), δ is equal to 0.17sec. Δ is set to 0.017sec considering 90% overlapping between consecutive windows. Therefore, the total number of windows over the whole epoch duration is *nWinds*=108 windows.

In this paper, we evaluated two popular modes of connectivity including phase-based metrics: (1) PLV (phase-locking value) and (2) PLI (phase-lag index), and one amplitude-based metric: (3) AEC (amplitude envelope correlation), as described below.

#### 4.1. Phase-locking value (PLV)

For each trial, the phase-locking value (Lachaux et al., 2000) that characterizes the phase relationship between two signals *x*(*t*) and *y*(*t*) is expressed as follows:

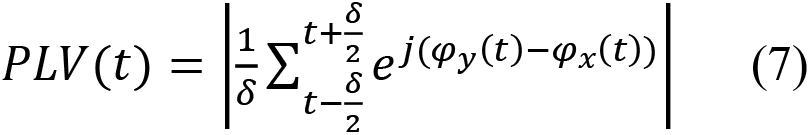

Where *φ_y_*(*t*) and *φ_x_*(*t*) represent the instantaneous phases of signals y and x respectively derived from the Hilbert transform at time window t. The Matlab function used to calculate PLV following Equation (6) is available on GitHub (https://github.com/judytabbal/dynCOALIA).

#### 4.2. Weighted phase-lag index (wPLI)

Unlike PLV, the weighted phase-lag index (wPLI) is insensitive to zero-lag interaction (Vinck et al., 2011). This index is based only on the imaginary component of the cross-spectrum and is thus robust to noise (Peraza et al., 2012).

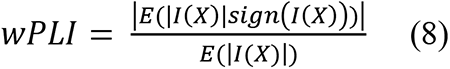

Where I(X) denotes the imaginary part of the signal’s cross-spectrum. Here, we used the Fieldtrip toolbox to compute wPLI (multi-taper method, fast Fourier transform, single Hanning tapper, 2Hz frequency resolution). As a certain amount of averaging across trials is required, wPLI was calculated at each temporal window using all trials of interest for each subject.

#### 4.3. Amplitude Envelope Correlation (AEC)

The amplitude envelope correlation (AEC) is calculated using the Hilbert transform of regional time series. Pearson correlation is then computed between the amplitude envelopes of two pairs of regions (Brookes et al., 2016; Hipp et al., 2012) for each trial. Refer to (https://github.com/judytabbal/dynCOALIA) for Matlab code implementation.

As a result, the output dimension of the dFC tensor was [*nROIs* × *nROIs* × *nWinds*] for every trial/subject. This tensor was unfolded into a 2D matrix 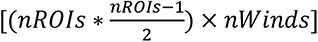 by removing redundant connections due to symmetry, followed by mean row subtraction. Finally, all trials/subjects dFC matrices were concatenated along temporal dimension and a group dFC matrix is constructed denoted ‘*P*’.

### 5. Dimensionality Reduction

This step is crucial to extract task-related brain network states (BNSs). It consists of summarizing all time-varying connectivity features in the constructed matrix ‘*P*’ 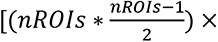 (*nWinds*Trials*nSubs*) into *k* dominant brain patterns over given time intervals. This problem can be formulated as follows:

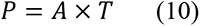

where ‘*A*’ [*nROIs*×*k*] is the mixing matrix illustrating the *k* spatial maps of dominant brain networks and ‘*T*’ [*k*×(*nWinds*\**Trials*\**nSubs*)] represents the corresponding temporal source signatures.

Among existing dimensionality reduction algorithms, we chose to investigate: (1) temporal Independent Component Analysis (tICA), (2) Principal Component Analysis (PCA), (3) Non-negative Matrix Factorization (NMF), and (4) Kmeans. They mainly differ in the constraints imposed on decomposed components. To reduce the effect of the number of states per method, we imposed *k*=6 components equal to the number of simulated networks in this work for all algorithms. We discuss this issue in the discussion section.

#### 5.1. temporal Independent Component Analysis (tICA)

tICA approach is already used by several studies (O’Neill et al., 2017; Yaesoubi et al., 2015) to derive *k* brain states that are ‘statistically mutually independent’ in time. Here, we evaluated tICA using three prominent ICA sub-methods: (1) JADE (Rutledge and Jouan-Rimbaud Bouveresse, 2013), (2) FastICA (Langlois et al., 2010), and (3) PSAUD (Becker et al., 2017). Briefly, FastICA is based on information theory, while JADE and PSAUD tend to optimize contrast functions based on high statistical order cumulants of the data. Technically, we adopted the following functions; ‘jader’ for JADE (Cardoso, 1999), ‘icasso’ for FastICA (Himberg and Hyvarinen, 2003), and ‘P_SAUD’ for PSAUD (Becker et al., 2017), implemented in Matlab (The Mathworks, USA, version 2019a).

#### 5.2. Principal Component Analysis (PCA)

PCA is a widely used technique that tends to reduce data dimensionality through a variance maximization approach. Hence, *k* orthogonal variables called ‘eigenvectors’ are extracted from a set of possibly correlated variables. Here, we applied the Singular Value Decomposition (SVD) algorithm of PCA (Golub and Reinsch, 1970) implemented in Matlab.

#### 5.3. Non-Negative Matrix Factorization (NMF)

Nonnegative matrix factorization (NMF) is an unsupervised machine-learning technique (Lee and Seung, 1999) that imposes a ‘positivity’ constraint on the decomposed factors. In this work, we selected the Alternating Least Squares (ALS) algorithm that has previously shown reliable performance in the fMRI context (Ding et al., 2013) with 100 times replications. For this purpose, the ‘nnmf’ Matlab function was used (Berry et al., 2007).

#### 5.4. Kmeans Clustering

Kmeans is one of the simplest clustering approaches (Lloyd, 1982). Based on feature similarity, Kmeans assigns each time point to one of the *k* centroids clusters, and the frequency of occurrence of each cluster at each time window is then calculated across all trials/subjects (Allen et al., 2014). In this study, we used L1 distance with 100 times replications and random centroid position initialization. We adopted the ‘kmeans’ function incorporated in Matlab (Mucha, 1986).

### 6. Performance Analysis

We estimate the similarity between reconstructed and reference sources/networks for each method. To this end, we defined four metrics: the ‘precision’, the ‘spatial similarity’, the ‘temporal similarity’, and the ‘global similarity’, to quantify the performance at each step of the pipeline.

#### 6.1. Inverse Model evaluation

The ‘precision’ metric was used to quantify the performance of inverse models at the cortical level. ‘Precision’ was calculated as the number of ‘correct’ regions divided by the total number of ‘active’ regions. In order to ensure that the density of estimated sources matches that of the ground truth that is necessary for the correct assessment of the precision value, we used a proportional threshold to keep the top *x*% regions with the highest source weights values. As we can see from the simulation scenario (Figure 2), the maximal number of simultaneously activated regions at a time interval is 7 corresponding to approximately *x* = 10% of the total region number (66 Desikan). In this work, we chose to vary the threshold from 10% to 15% for ‘precision’ calculation, and then take the average value across threshold values.

#### 6.2. Inverse Model/Functional connectivity combination evaluation

The ‘spatial similarity’ metric was used to quantify the performance of different inverse model/functional connectivity combination methods at the network level. Briefly, this measure takes into consideration the distribution of weights across edges between and within brain lobes. In particular, the brain is decomposed into 7 main lobes (occipital, parietal, temporal, central, frontal, prefrontal, limbic) yielding 28 possible connection types (i.e., occipital-parietal, occipital-temporal, central-prefrontal, temporal-temporal…). Then, the accuracy of connectivity weights between reference and reconstructed networks is calculated for each connection type. The spatial similarity is finally defined as the averaged value of accuracy over all connection types. The reader can refer to supplementary materials (Figure S2) for a more detailed description of spatial similarity calculation. The proportional threshold was also used to retain the top-weighted edges, varied from top 1% to 2% of total undirected possible edges, which correspond to the possible number of connections between the selected range of nodes density (see section 6.1).

#### 6.3. Decomposition methods evaluation

For each dimensionality reduction method, the ‘temporal similarity’ (correlation between the reference and reconstructed temporal signals) and ‘spatial similarity’ (explained in the previous section) were computed between the reference and the ‘best-matched’ components to evaluate both the temporal and spatial performance of each method. Then, ‘global similarity’ was calculated as the average between both similarities.

### 7. Statistical Analysis

We also applied our pipeline with performance analysis at subject level data yielding a distribution set of values across 20 subjects. Then, the ANOVA test was used (The Mathworks, USA, version 2019a) to statistically quantify differences between the tested methods, followed by a post-hoc correction for multiple comparisons (using the built-in function ‘multcomp’) with Bonferroni correction (statistical significance *p*-value=0.01).

## Data availability

All the simulated data supporting the findings of this study are available on the Github (https://github.com/judytabbal/dynCOALIA).

## Code availability

All codes supporting the results of this paper can be found at (https://github.com/judytabbal/dynCOALIA). All analysis codes were implemented and performed in Matlab software using several toolboxes such as Fieldtrip (for BEM, eLORETA, and wPLI computation), EEGLAB (for JADE computation), and other hand-written/customized Matlab scripts and functions.

## Results

### 1. Source localization: wMNE vs. eLORETA

To first visualize results at the cortical level, we illustrate reconstructed brain sources in Figure 3 at simulated time intervals for each inverse model (wMNE and eLORETA) along with both source space resolution (LowRes vs. HighRes), where ‘LowRes’ denotes the lead field calculation on 66 Desikan regions directly while ‘HighRes’ refers to lead field calculation on high-resolution cortex followed by projection (average) on 66 Desikan regions (see Materials and Methods). Results shown were averaged over trials and subjects.

**Figure 3.**
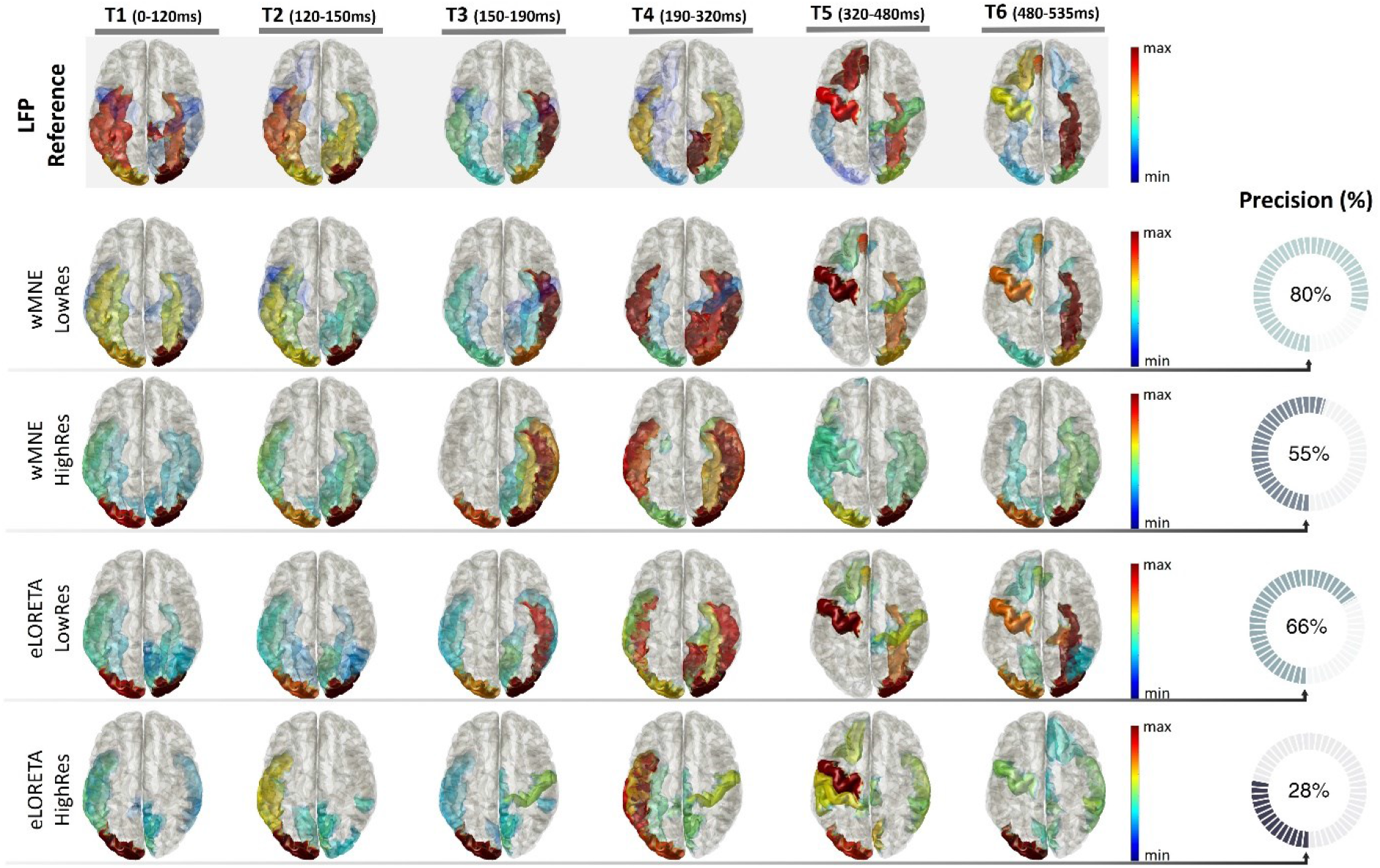
Inverse model evaluation (wMNE and eLORETA) using different source space resolutions. LowRes refers to lead field computation on 66 Desikan regions and HighRes refers to lead field computation on high-resolution cortex 15002 followed by projection on 66 Desikan regions. Reconstructed brain sources are shown at the group level (averaged over trials and subjects). The top 10 activated regions are visualized for each method and at each time interval. Brain regions’ colors are based on their weights (blue for the less activated, red for the most activated). The precision value was calculated relative to LFP reference sources at each time interval with a proportional threshold ranging from 10% to 15% of the total node number. The precision value is shown on the right side of each method as the average value over all time intervals and threshold values.

For inverse methods quantification, we calculated the precision metric averaged over subjects’ data (see Materials and Methods) at each simulated time interval between each method and reference sources. In our case, reference sources correspond to the simulated LFP cortical-level data generated by the COALIA model (as illustrated in the first row of Figure 3). Then, precision values were averaged over time intervals to obtain a single quantification value for each method. wMNE showed 80% precision with LowRes and 55% with HighRes while eLORETA showed 66% with LowResres66 and 28% with HighRes (see Figure 3).

We can notice that (1) wMNE outperformed eLORETA for both resolutions and (2) the lead field computed directly on the regions of interest seems to lead to higher precision results for both inverse models.

### 2. Inverse solutions and connectivity measures combination

To visualize results at the network level, we illustrate in Figure 4 the estimated dynamic functional connectivity matrices at time intervals for each inverse model/functional connectivity method. For visualization purposes, results were averaged over trials and subjects. For results quantification, we calculated the spatial similarity metric (see supplementary Figure S2) at each time interval between each method and corresponding reference networks. In our case, reference networks correspond to the functional connectivity method applied to simulated LFP sources data. For example, brain networks estimated from PLV applied on LFP sources were considered as a reference for wMNE/PLV and eLORETA/PLV. We followed the same concept for other methods. Then, spatial similarity values were averaged over time intervals to obtain a single quantification value for each method. At the group-level, global spatial similarities were 0.65 for wMNE/PLV, 0.62 for wMNE/wPLI, 0.61 for wMNE/AEC, 0.53 for eLORETA/PLV, 0.52 for eLORETA/wPLI and 0.50 for eLORETA/AEC (see Figure 6(a)).

**Figure 4.**
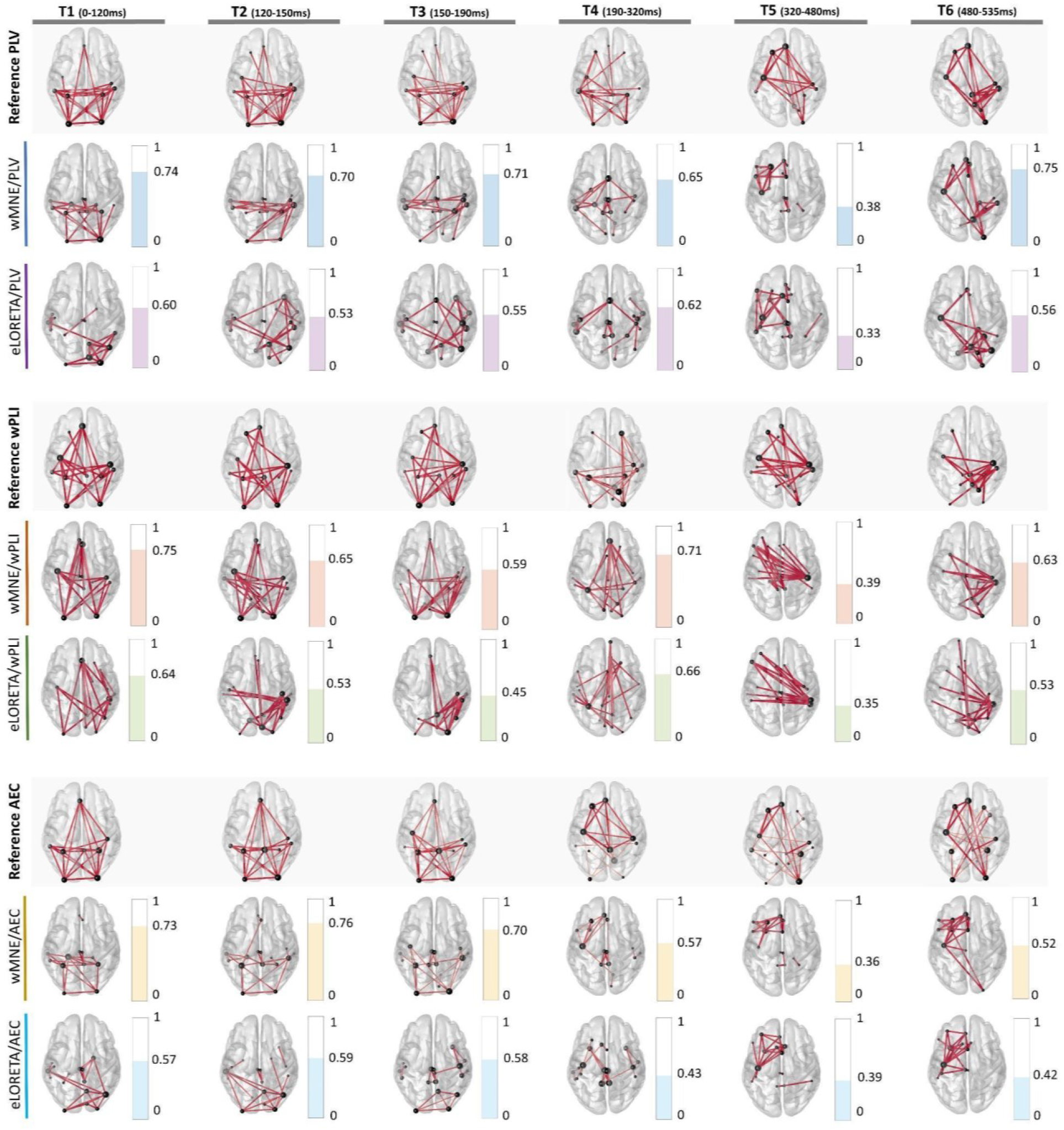
Inverse model/functional connectivity combination evaluation. Reconstructed brain networks are shown at the group level (averaged over trials and subjects). All brain networks were thresholded for visualization by keeping the top 1% edges with the highest connectivity values. Edges line width indicates connectivity strength and nodes sphere size reveals brain region strength. Spatial Similarity was calculated between reconstructed and reference brain networks at each time interval with a proportional threshold ranging from 1% to 2% of all possible connectivities. The spatial similarity value is indicated on the right side of each network as the average value over threshold values. PLV networks applied on LFP sources were considered as reference brain networks for wMNE/PLV and eLORETA/PLV. wPLI networks applied on LFP sources were considered as reference brain networks for wMNE/wPLI and eLORETA/wPLI. AEC networks applied on LFP sources were considered as reference brain networks for wMNE/AEC and eLORETA/AEC. A color code was attributed to each method (blue for wMNE/PLV, orange for wMNE/wPLI, yellow for wMNE/AEC, purple for eLORETA/PLV, green for eLORETA/wPLI, and cyan for eLORETA/AEC).

At the group level, one can notice that in the current simulation instance (1) the highest similarity was reached for wMNE/PLV while eLORETA/AEC performed the worst. (2) wMNE performed better than eLORETA for all functional connectivity measures and (3) PLV exhibited the highest similarity values followed by wPLI and then AEC for both source reconstruction methods. Estimated networks were also averaged over trials for each subject and similarity was computed at subject-level data. Similarity results of the distribution over 20 subjects are displayed on boxplots in Figure 6(c) Statistical analysis performed by ANOVA test showed non-significant differences between (1) wMNE/wPLI and wMNE/AEC (*p*>0.05) and (2) eLORETA/PLV and eLORETA/wPLI (*p*>0.05) while all other combinations showed significant differences (*p*<0.05).

### 3. Dimensionality reduction methods evaluation

As we are dealing with six simulated networks, we imposed ‘k=6’ states for each dimensionality reduction method. As a result, six dynamic brain states (denoted ‘S_i_’ in Figure 5), including spatial maps and corresponding temporal signals, were extracted from dynamic functional connectivity networks. Here, we chose estimated networks from the best combination of previously evaluated inverse model/functional connectivity methods (wMNE/PLV, with ‘LowRes’) (denoted ‘C_i_’ in Figure 5) to be considered as reference networks for spatial maps evaluation. The six simulated time intervals denoted ‘T_i_’ were considered as reference temporal occupancy for temporal signals evaluation. In this context, spatial similarity and temporal similarity were computed between each connectivity reference and all extracted states. Then, the global similarity was calculated as the average between both spatial and temporal similarities. Following this, each reference connectivity ‘C_i_’ was matched with the most representative state ‘S_j_’ having the highest global similarity value among all extracted states. Hereafter, we refer to the ‘maximal global similarity’ as the global similarity value between the reference state and the corresponding best-matched reconstructed state. For example, for PSAUD, results show that C_1_ matched the best S_1_ (maximal global similarity=0.88), C_2_ matched S_2_ (maximal global similarity=0.71), C_3_ matched S_2_ (maximal global similarity=0.68), C_4_ matched S_3_ (maximal global similarity=0.87), C_5_ matched S_4_ (maximal global similarity=0.75), C_6_ matched S_6_ (maximal global similarity=0.64). PSAUD results are expressed in Figure 5 and all other dimensionality reduction methods results are detailed in supplementary materials (Figure S3, S4).

**Figure 5.**
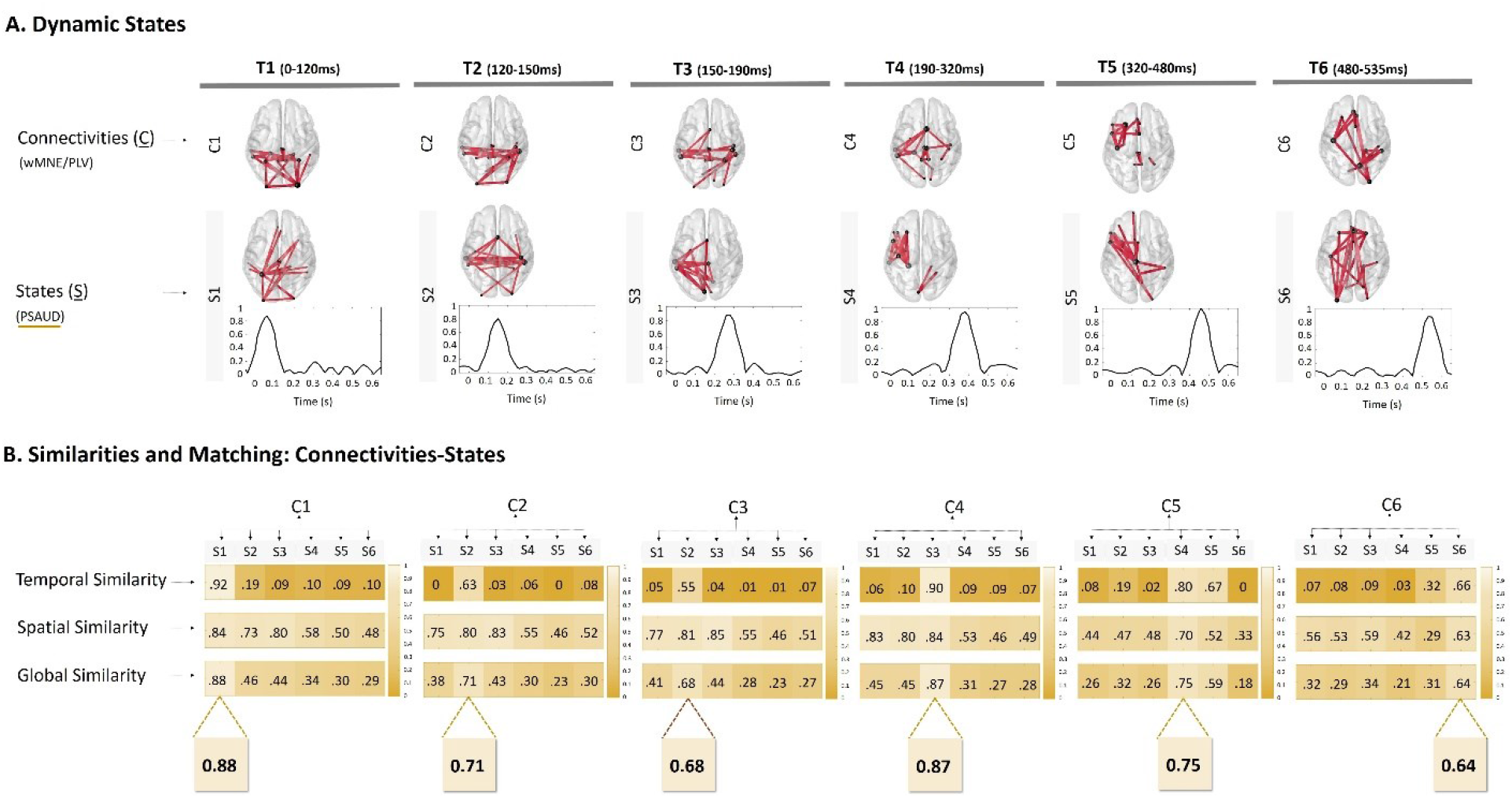
A typical example of the ICA (‘PSAUD’) method evaluation: is PSAUD’. Results of all other evaluated dimensionality reduction methods (JADE, FastICA, PCA, NMF, and Kmeans) are detailed in supplementary materials (Figure S3.). A. Connectivities denoted ‘Ci’ represent our reference components and correspond to wMNE/PLV networks calculated at each time interval. Dynamic States results denoted ‘Si’ (spatial maps with temporal activity) are shown at the group level for the ‘PSAUD’ method (number of states=6). All networks were thresholded for visualization purposes (top 1% edges with the highest connectivity values). B. Temporal and Spatial Similarities between all extracted states and each reference connectivity were computed. Then, the global similarity was calculated as the average value between spatial and temporal similarity. Similarity values are represented by different color shades (the higher the value, the brighter the color is). Each reference connectivity ‘Ci’ is then matched with the state ‘Sj’ that corresponds to the highest global similarity value (i.e. C1 matches S1 with a global similarity equal to 88%, C5 matches S4 with a global similarity equal to 75%…).

**Figure 6.**
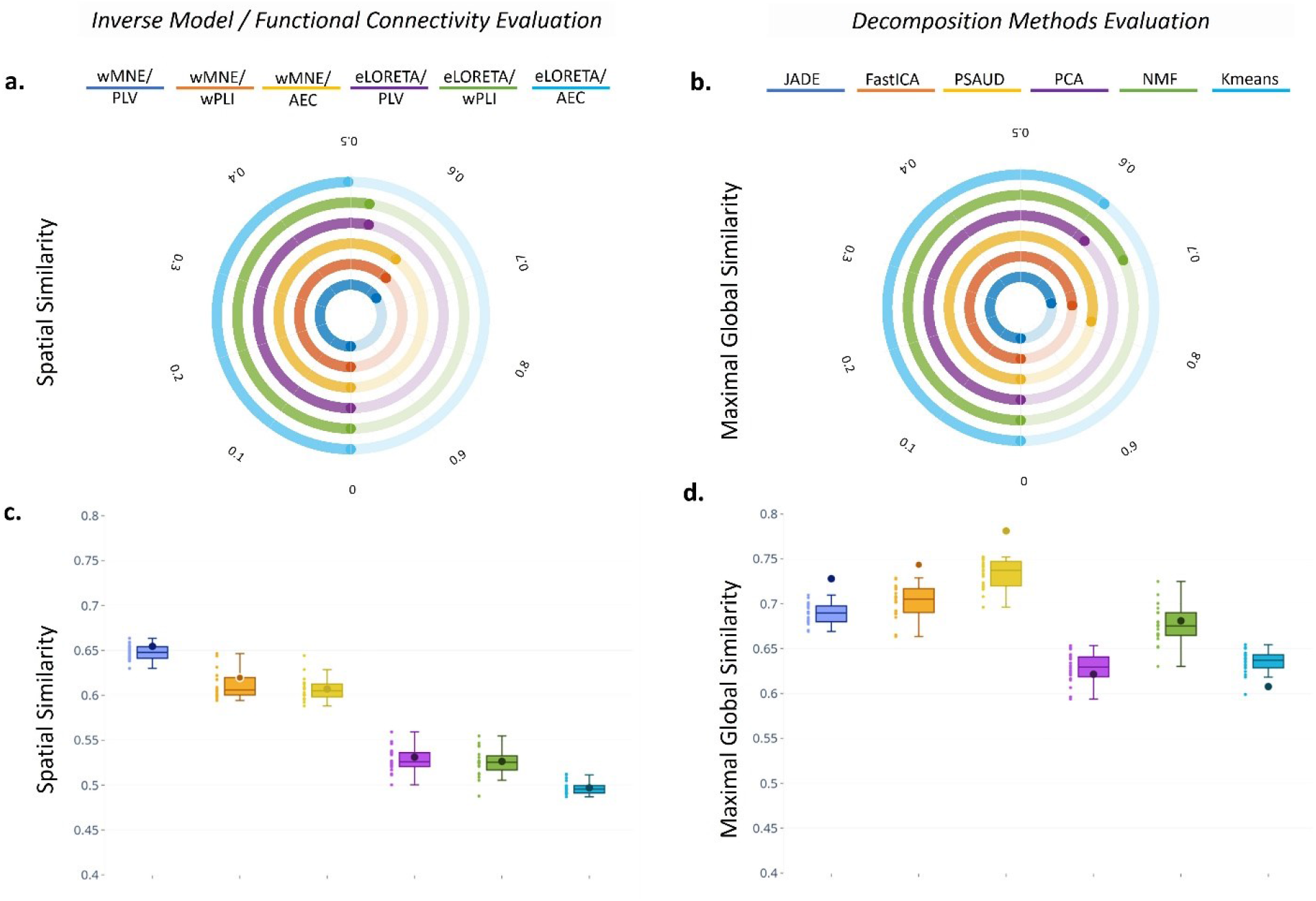
Group-level and Subject-level overall methodology evaluation. Spatial similarity averaged over all time intervals was used as an overall evaluation measure for different inverse model/functional connectivity combinations at the group-level (a.) and subject-level (c.). Maximal Global similarity averaged over all time intervals was used as an overall evaluation measure for different dimensionality reduction methods at the group-level (b.) and subject-level (d.). Each method is represented by a specific color. Circular bars and boxplots were used to visualize results at both group and subject levels respectively. The results of each subject are represented by small circles on the left side of each boxplot. We also overlaid results obtained in (a) and (b) in the boxplots in (c) and (d) respectively as centered bold circles to show subject-level results relative to the group level.

To globally quantify each method, we averaged maximal global similarity values obtained across all time intervals. At the group-level, maximal global similarity values were 0.73 for JADE, 0.74 for FastICA, 0.78 for PSAUD, 0.62 for PCA, 0.68 for NMF and 0.61 for Kmeans (see Figure 6(b)).

Dimensionality reduction methods were also tested at subject-level data where all trial dFC matrices were concatenated along temporal dimensions for each subject. Maximal Global Similarity results distribution over 20 subjects are displayed on boxplots in Figure 6(d) (mean values: 0.69 for JADE, 0.70 for FastICA, 0.73 for PSAUD, 0.63 for PCA, 0.68 for NMF and 0.63 for Kmeans). Statistical analysis performed by ANOVA test showed non-significant differences between (1) JADE and FastICA (p>0.05), (2) JADE and NMF, and (3) PCA and Kmeans (p>0.05) while all other combinations showed significant differences (p<0.05).

Similarly, we evaluated separately the Spatial and Temporal Similarity distribution over 20 subjects of the best-matched states, as shown in the boxplots of Figure S5 in Supplementary Materials, followed by the statistical ANOVA test. We aim to deepen the performance examination of our tested methods to consider each of the spatial and temporal modes. For the spatial similarity, there were no significant differences (p-value>0.05) between PSAUD (mean=0.7371), NMF (mean=0.7269) and Kmeans (mean=0.7357), having the highest values among all methods, followed by FastICA (mean=0.6955) and PCA (mean=0.6783) with non-significant differences, then JADE (mean=0.6419). On the other hand, PSAUD (mean=0.7294) exhibited similar temporal performance to each of JADE (mean=0.7355) and FastICA (mean=0.7088) (p-value>0.05). Weaker temporal similarities (with significant differences) were shown for NMF (mean=0.6257), PCA (mean=0.5787), and Kmeans (mean=0.5343).

Moreover, we tested dimensionality reduction methods’ dependency on measurement noise (see Figure S6). The maximal global similarity is plotted against different noise levels expressed by the λ value for JADE (*mean* ± *std* = 0.68±0.0216), FastICA (*mean* ± *std* = 0.72±0.0060), PSAUD (*mean* ± *std* = 0.73±0.0228), PCA (*mean* ± *std* = 0.64±0.0212), NMF (*mean* ± *std* = 0.68±0.0255), Kmeans (*mean* ± *std* = 0.64±0.0276). Results were slightly influenced by noise with Kmeans being relatively the most dependable method (with the highest std value) and FastICA as the least dependent method.

## Discussion

Tracking dynamic brain networks from non-invasive electrophysiological (EEG/MEG) data is a key challenge in neuroscience. A critical issue is to evaluate to what extent extracted ‘brain network states’ match those that are truly activated during tasks. In this context, the presence of ‘ground truth’ data is of utmost importance to ensure objective and quantitative evaluation of pipeline processes. The main contribution of this study is the use of simulated EEG data - in a dynamic context - from a physiologically grounded computational model as ground truth for evaluating methods’ performance in ‘re-estimating’ correctly reference states. The dynamic approach was simulated by adopting the scenario of a picture-naming task evolving several brain states fluctuating over time (Hassan et al., 2015; Mheich et al., 2021, 2015).

### Methods evaluation

In this paper, we conducted a systematic evaluation analysis of two source reconstruction methods (wMNE/eLORETA), three functional connectivity measures (PLV/wPLI/AEC), and six dimensionality reduction methods (JADE/FastICA/PSAUD/PCA/NMF/Kmeans) used as a three-step pathway to extracting dynamic brain network states from scalp-EEG signals. Overall, results show that wMNE outperforms eLORETA in the context of our simulation study. This is observed when evaluating source reconstruction methods on both cortical and network levels. In this study, the wMNE/PLV combination exhibits the best performance among all inverse model/functional connectivity combinations tested. Interestingly, these results are in line with previous comparative studies showing consistency and robust results for wMNE/PLV combination in the context of EEG source space connectivity using real data from picture naming task (Hassan et al., 2014) and simulated data from epileptogenic-modeled networks (Allouch et al., 2020; Hassan et al., 2017). The strength of PLV results may reflect the potential mechanisms of the existing zero-lag synchronization of neural activity as already discussed in previous works (Gollo et al., 2014; Roelfsema et al., 1997). It is worth noting that, in the particular context of the study, functional connectivity methods had less impact on results than inverse models. For instance, wPLI-AEC performed similarly with wMNE, and PLV-wPLI performed similarly with eLORETA.

We also explored a technical point related to the source space resolution used for lead field computation. In this context, there is no clear evidence about the ideal source grid resolution to adopt. Our results suggest that computing the lead field after cortex parcellation (referred to as LowRes) provides higher accuracy results than lead field computation before parcellation followed by ROI projection (referred to as HighRes). One explanation might be related to the induced blurring effect between close sources when using a high-resolution source grid, which might increase the cross-talk between specific cortical locations (Maldjian et al., 2014; Schoffelen and Gross, 2009).

Overall, performance analysis showed promising results for most dimensionality reduction methods regarding their ability to derive dominant brain states. In this work, global similarity results highlight the performance of ICA subtypes methods relative to other methods. These findings are in line with a recent comparative study between source separation methods applied to three independent real MEG datasets (Tabbal et al., 2021). Particularly, in this work, it is noteworthy to point out that PCA, NMF, and Kmeans exhibit fragility in the temporal domain reflected by relatively low temporal similarities at several intervals (Figure S3). This is also reflected by the statistical comparative analysis between the evaluated methods at the level of the spatial and temporal domain (Figure S5). For instance, although NMF and Kmeans were able to extract the correct spatial networks with high precision relative to other techniques, these methods lack the potential to accurately track their fast temporal activity, which notably influenced the global methods’ performance (Figure 6). In contrast, ICA techniques were significantly more sensitive than other techniques to the temporal fluctuations of the simulated networks. We can notice an attractive behavior for the PSAUD method in terms of spatial and temporal precision. Therefore, when studying the dynamics of task-related brain activity, it is important to choose a decomposition technique that assures good spatial and temporal accuracy in the functional brain networks.

Furthermore, it is crucial to note that most methods revealed weaker performance at specific time intervals (T2, T3, and T6) (Figure S4). Interestingly, these intervals are the narrowest (T2 lasts for 30ms, T3: 40ms, and T6: 55ms) compared to other simulated intervals (T1: 120ms, T4: 130ms, and T5: 160ms). This point is of particular interest as it shows method limitation when applied on a very fast timescale (∼<100ms). The noise level was varied to test the effect of scalp-level noise on extracted brain states. Although most dimensionality reduction methods were robust against noise variation, a generalized stability assessment of each method relative to measurement noise needs more detailed investigation related to the type and level of the realistic added noise.

Finally, the subject-level and group-level results seem to be convergent (Figure 6(c), 6(d)). We believe that these results are of significant neuroscientific interest, in particular, for application in clinical neuroscience, which needs consistent and reliable results at a patient level. Therefore, it can open new avenues for detailed methodology/parameters evaluation and optimization of this framework at the single-subject data.

### Methodological Considerations

In this study, we aim to carry out a dynamic analysis of brain network activity using a computational model serving as a ground truth for our pipeline methodology evaluation. To this end, one could define any set of brain network states as input to the COALIA model. Here, simulated states were determined based on a realistic picture naming task scenario, for which a solid background is available concerning activated brain regions and networks. Furthermore, the variety of brain regions activated distinctly from the onset to the reaction time is a major benefit of this scenario choice as we are interested in evaluating the dynamic behavior of methods at a realistic and rapid timescale. However, it is noteworthy to mention that the current study is an exemplar-based analysis for methods performance. Hence, results generalization requires more exhaustive evaluation over many simulation instances where other scenarios could be also tested simulating other simple or complex tasks or even customized scenarios including pre-defined brain states with manipulation in time length, number of activated regions, and states.

Besides, one limitation in this work is related to the high signal-to-noise ratio of the simulated brain sources data relative to the real electrophysiological data. Indeed, further investigations about the type/value of the noise added to the simulated gamma oscillations are needed to reduce, as much as possible, the gap between our brain model and the real brain physiology.

The main objective of this paper is to assess the ability of well-known existing methodologies to track dominant brain network states from EEG signals, rather than to perform an exhaustive comparative analysis of all possible combinations of the ‘three-step’ pipeline methods. Nevertheless, our analysis work could be extended to cover a much more variety of techniques at each step. First, beamformer family methods (Van Veen et al., 1997) could be evaluated besides the minimum norm estimates methods tested in this paper. Second, as we accounted for the volume conduction effect in phase-based coupling measures by keeping and removing zero-lag synchronization using PLV and wPLI methods, it would be also interesting to evaluate the effect of leakage correction on amplitude-based coupling measures. Therefore, we suggest adding the corrected version of amplitude envelope correlation (AEC-c) to the comparative analysis ultimately. Furthermore, we adopted a sliding window approach to study the dynamic behavior of functional connectivity. Although window size was calculated corresponding to the minimal length required (Lachaux et al., 2000), the generalization of this finding to amplitude-based functional connectivity metrics may be critical, and therefore, further efforts are required to ensure a precise evaluation. Moreover, several findings criticize the performance of sliding windows when computed over very short epoch durations (Fraschini et al., 2016; Liuzzi et al., 2019). We, therefore, suggest testing high-resolution measures of functional connectivity that are sensitive to fast fluctuations (Tewarie et al., 2019) as an extension to our simulation study.

Third, to extract dominant brain states, we tested decomposition methods including source separation and clustering techniques. Nevertheless, it will be interesting to add other strategies that revealed accurate results in previous studies as Hidden Markov Model (HMM) (Vidaurre et al., 2018) and the multivariate autoregressive model (Casorso et al., 2019).

The connectivity matrices were thresholded using proportional thresholding to standardize our comparison across methods, trials, and intervals. Although we accounted for the effect of the threshold value, one limitation is related to the fact that a priori knowledge about network density is not available in real experimental data contrary to simulated data. Thus, in the context of real data, the selection of the appropriate threshold values is critical and crucial for further investigation, which is out of the scope of this paper.

In addition, the number of reference states is known a priori in the simulation-based analysis, but often hard to predict in the case of empirical data. The optimal number of brain network states could be defined based on several optimization criteria, such as the cross-validation criterion (Pascual-Marqui et al., 1995), Krzanowski-Lai criterion (Krzanowski and Lai, 1988), kneedle algorithm (Satopaa et al., 2011), and difference of data fitting (DIFFIT) (Timmerman and Kiers, 2000; Wang et al., 2018). However, to keep our analysis compact, we decided to fix this parameter to the exact number of simulated states (6 states) for all decomposition methods. Although this could pose a limitation concerning the optimum performance of each specific algorithm, the focus of the present work is the direct comparative evaluation of different algorithms based on pre-defined reference networks. For instance, applying a specific optimization criterion to simultaneously all algorithms may fit better some algorithms than others, and hence, influence the interpretation of our results. Therefore, future studies may test different existing approaches used to optimize the number of derived components relative to each algorithm.

Finally, in this work, we set the number of EEG channels (257 channels) since it established accurate source localization results relative to lower sensor densities (Allouch et al., 2020; Song et al., 2015). It can be however interesting to follow the work of such studies and examine the effect of sensor spatial resolution by decreasing successively the number of electrodes from high-density (257 channels) to low-density (19 channels) EEG signals.

## Conclusion

In this study, we propose a complete framework for analyzing the task-related dynamic electrophysiological brain networks. We used a physiologically inspired full-brain model (named COALIA) as ‘ground-truth’ to systematically validate and optimize the pipeline study including EEG-source space connectivity estimation and dynamic brain network states extraction. As a proof of concept, the study was conducted in a dynamic scenario that emulates the picture naming task. Our findings suggest a good performance of the wMNE/PLV combination to elucidate the appropriate functional networks. Results also revealed the promising efficiency of ICA techniques to derive relevant dynamic brain network states. Using this framework, researchers are invited to generate other tasks for further validation and address additional methodological points involved in the optimization of the EEG/MEG source-space network analysis.

