## Supplementary Materials for "Assessing HD-EEG functional connectivity states using a human brain computational model"

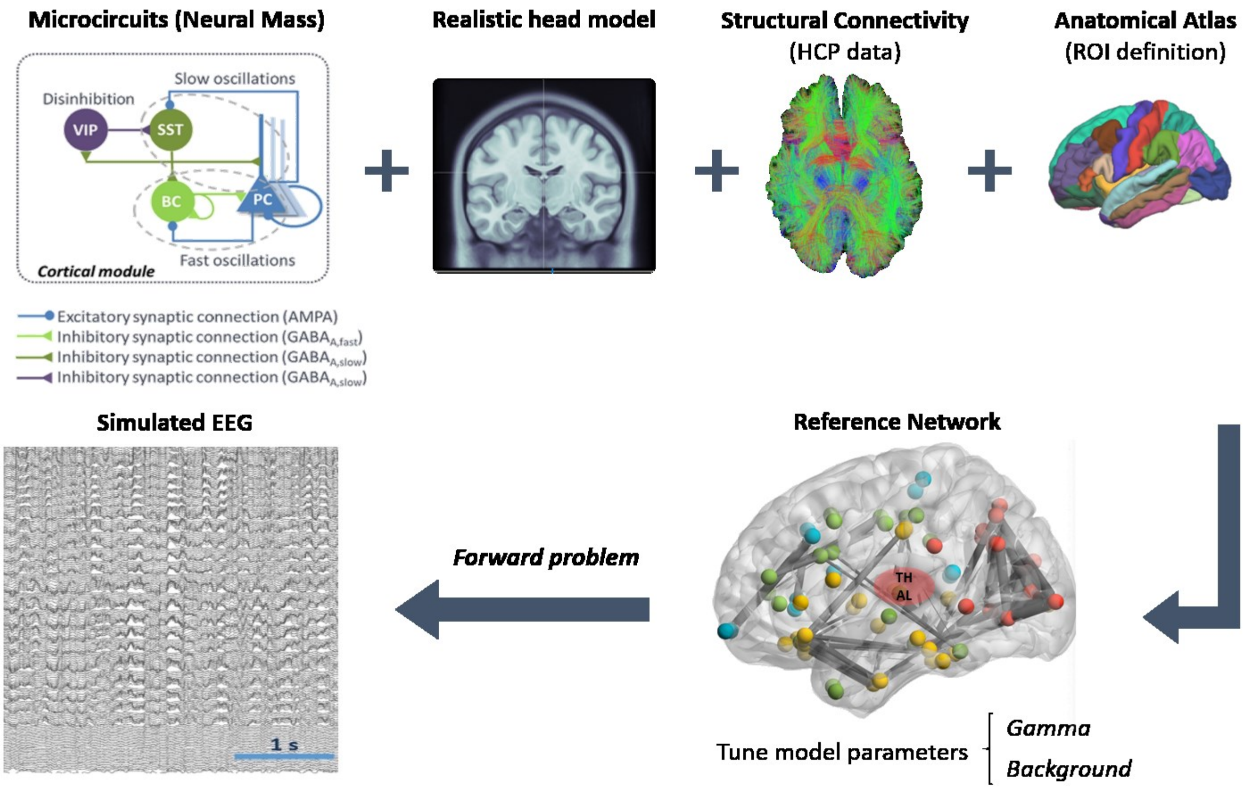


*Figure S1. Overview description of the updated COALIA model used in our study to simulate EEG signals. At the local level, the model is formed by microcircuits based on neural mass. Each neural mass consists of subsets of excitatory and inhibitory neural population. At the global level, the cortical-level reference network is defined by taking into account several parameters including realistic head model, structural connectivity from Human Connectome Project (HCP) and anatomical desikan atlas to define regions of interest. Finally, the simulated EEG data are computed by solving the forward model.*

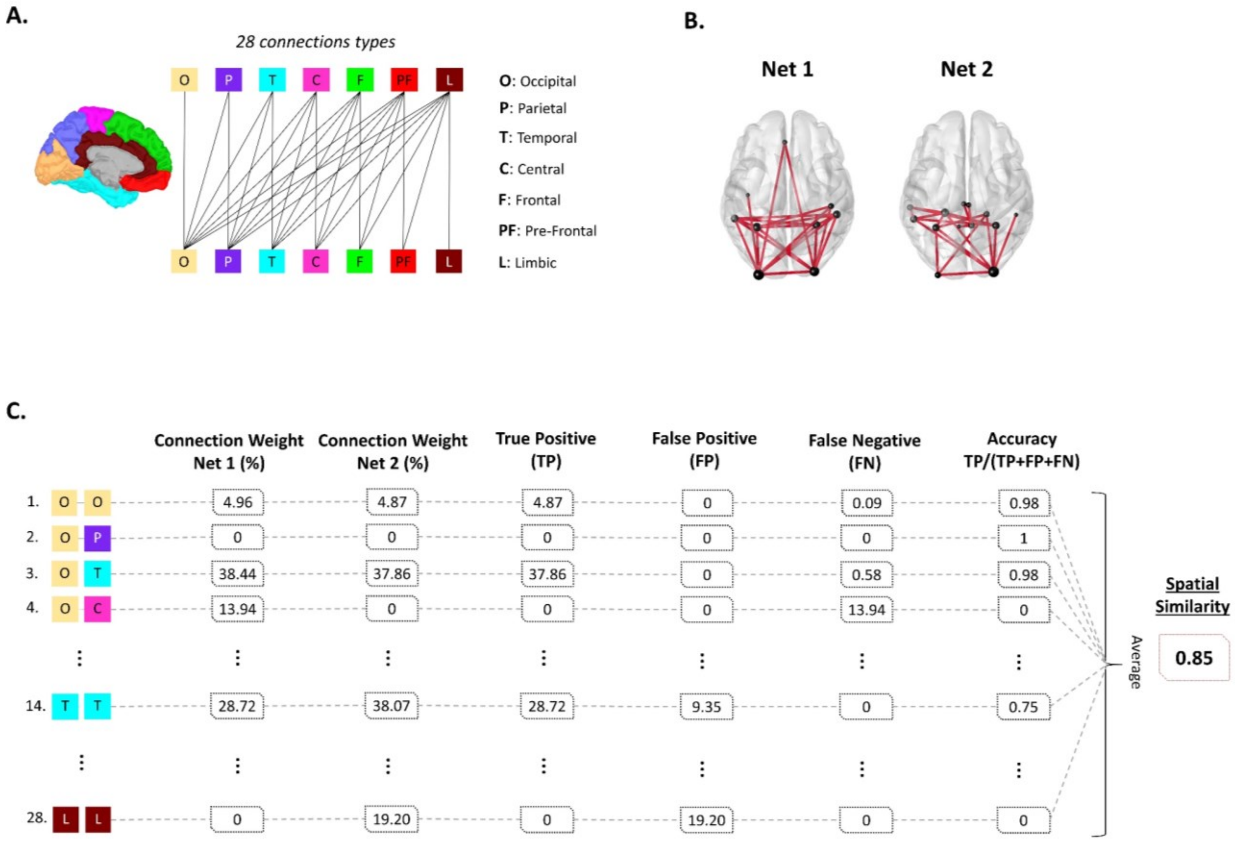


*Figure S2. (A) Lobes distribution on the brain cortex used for spatial similarity calculation. The brain cortex is divided into seven main lobes (Brainstorm distribution) including occipital, parietal, temporal, central, frontal, prefrontal and limbic. In total, 28 possible connections types can be established between different brain lobes (i.e. O-O refers to the existing connections between nodes of occipital lobe, while P-T refers to existing connections between nodes in parietal lobe and nodes in temporal lobe). (B) Two exemplar brain networks to show spatial similarity calculation. Net 1 refers to reference network, Net 2 refers to reconstructed network. (C) Detailed description of spatial similarity calculation procedure. For each connection type, connection weights of both networks are computed. True Positive (TP) represents common connection weight between both networks., False Positive (FP) represents connection weight present in Net 2 exclusively. False Negative (FN) represents connection weight present in Net 1 exclusively. Spatial similarity between networks is finally calculated as the average of accuracy over all connections types.*

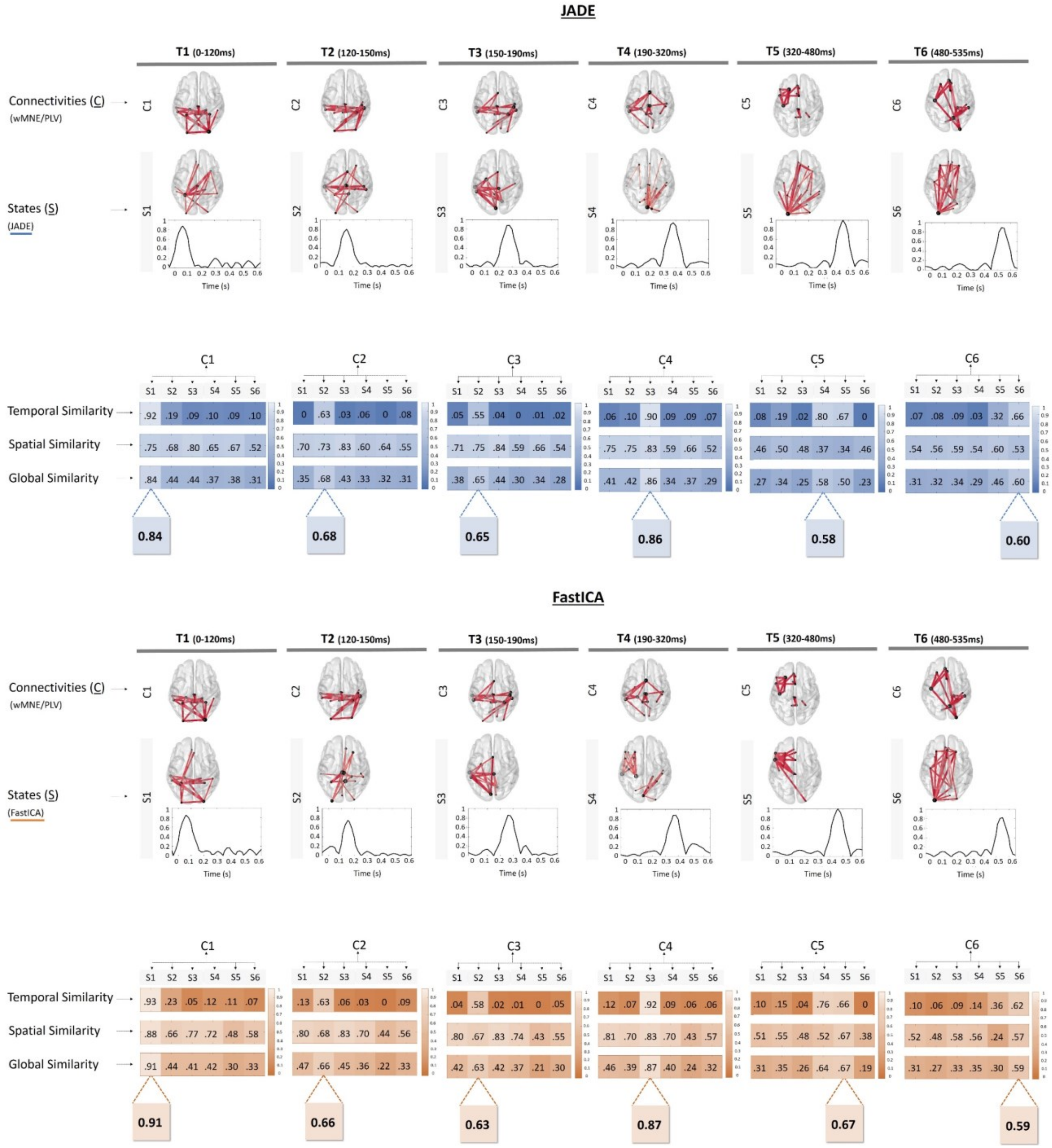
JADE
T1 (0-120ms)
T2 (120-150ms)
T3 (150-190ms)
T4 (190-320ms)
T5 (320-480ms)
T6 (480-535ms)
Connectivities (C)
(wMNE/PLV)
C1 C2 C3 C4 C5 C6
States (S)
(JADE)
S1 S2 S3 S4 S5 S6
Time (s)
Temporal Similarity
Spatial Similarity
Global Similarity
C1: .92 .19 .09 .10 .09 .10 / .75 .68 .80 .65 .67 .52 / .84 .44 .44 .37 .38 .31
C2: 0 .63 .03 .06 0 .08 / .70 .73 .83 .60 .64 .55 / .35 .68 .43 .33 .32 .31
C3: .05 .55 .04 0 .01 .02 / .71 .75 .84 .59 .66 .54 / .38 .65 .44 .30 .34 .28
C4: .06 .10 .90 .09 .09 .07 / .75 .75 .83 .59 .66 .52 / .41 .42 .86 .34 .37 .29
C5: .08 .19 .02 .80 .67 0 / .46 .50 .48 .37 .34 .46 / .27 .34 .25 .58 .50 .23
C6: .07 .08 .09 .03 .32 .66 / .54 .56 .59 .54 .60 .53 / .31 .32 .34 .29 .46 .60
0.84
0.68
0.65
0.86
0.58
0.60
FastICA
T1 (0-120ms)
T2 (120-150ms)
T3 (150-190ms)
T4 (190-320ms)
T5 (320-480ms)
T6 (480-535ms)
Connectivities (C)
(wMNE/PLV)
States (S)
(FastICA)
C1: .93 .23 .05 .12 .11 .07 / .88 .66 .77 .72 .48 .58 / .91 .44 .41 .42 .30 .33
C2: .13 .63 .06 .03 0 .09 / .80 .68 .83 .70 .44 .56 / .47 .66 .45 .36 .22 .33
C3: .04 .58 .02 .01 0 .05 / .80 .67 .83 .74 .43 .55 / .42 .63 .42 .37 .21 .30
C4: .12 .07 .92 .09 .06 .06 / .81 .70 .83 .70 .43 .57 / .46 .39 .87 .40 .24 .32
C5: .10 .15 .04 .76 .66 0 / .51 .55 .48 .52 .67 .38 / .31 .35 .26 .64 .67 .19
C6: .10 .06 .09 .14 .36 .62 / .52 .48 .58 .56 .24 .57 / .31 .27 .33 .35 .30 .59
0.91
0.66
0.63
0.87
0.67
0.59

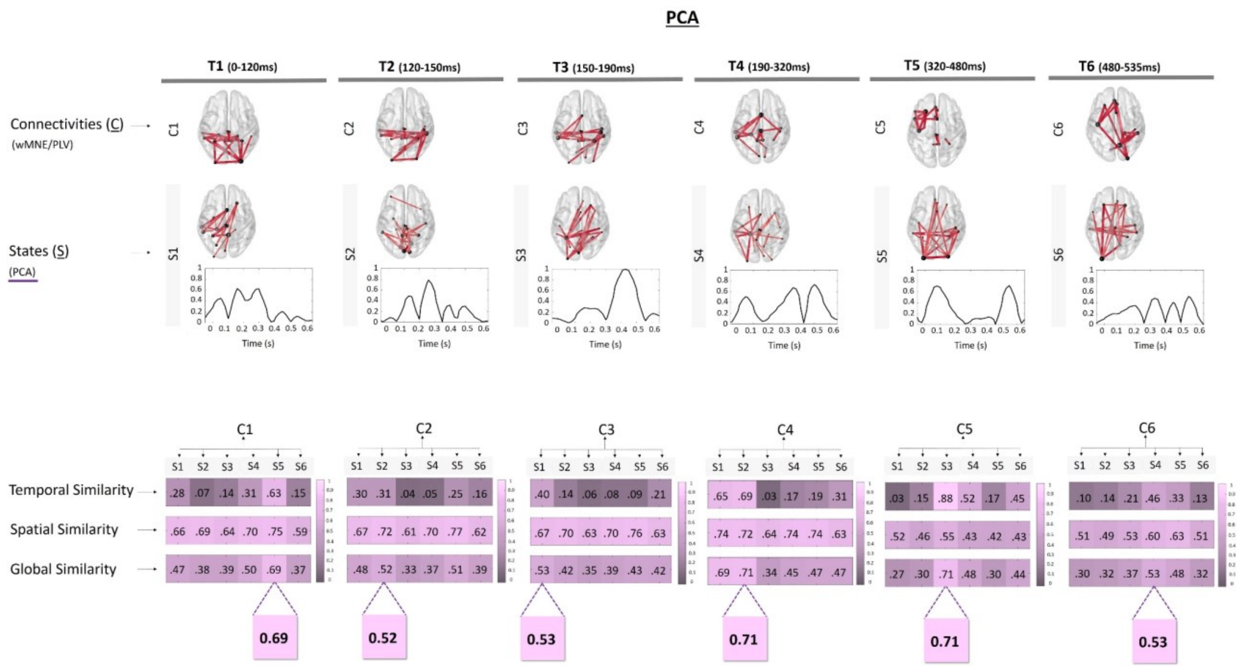
PCA
T1 (0-120ms)
T2 (120-150ms)
T3 (150-190ms)
T4 (190-320ms)
T5 (320-480ms)
T6 (480-535ms)
Connectivities (C)
(wMNE/PLV)
States (S)
(PCA)
Time (s)
Temporal Similarity
Spatial Similarity
Global Similarity
C1: S1 S2 S3 S4 S5 S6
.28 .07 .14 .31 .63 .15
.66 .69 .64 .70 .75 .59
.47 .38 .39 .50 .69 .37
0.69
C2: S1 S2 S3 S4 S5 S6
.30 .31 .04 .05 .25 .16
.67 .72 .61 .70 .77 .62
.48 .52 .33 .37 .51 .39
0.52
C3: S1 S2 S3 S4 S5 S6
.40 .14 .06 .08 .09 .21
.67 .70 .63 .70 .76 .63
.53 .42 .35 .39 .43 .42
0.53
C4: S1 S2 S3 S4 S5 S6
.65 .69 .03 .17 .19 .31
.74 .72 .64 .74 .74 .63
.69 .71 .34 .45 .47 .47
0.71
C5: S1 S2 S3 S4 S5 S6
.03 .15 .88 .52 .17 .45
.52 .46 .55 .43 .42 .43
.27 .30 .71 .48 .30 .44
0.71
C6: S1 S2 S3 S4 S5 S6
.10 .14 .21 .46 .33 .13
.51 .49 .53 .60 .63 .51
.30 .32 .37 .53 .48 .32
0.53

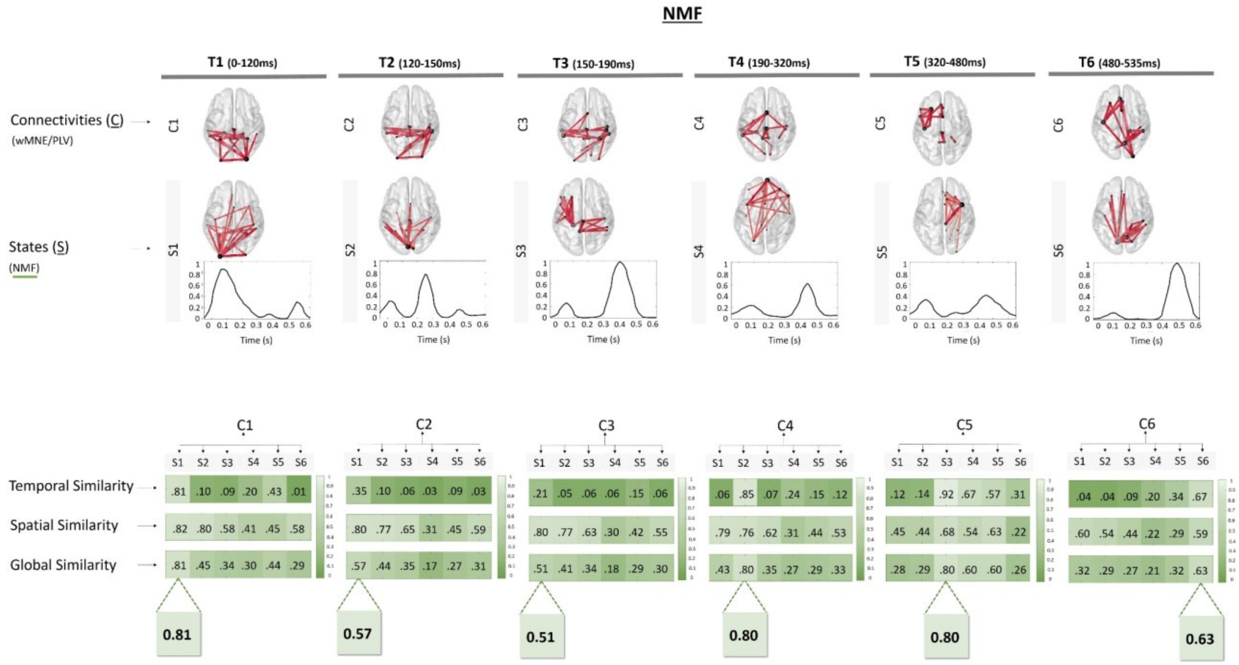
NMF
T1 (0-120ms)
T2 (120-150ms)
T3 (150-190ms)
T4 (190-320ms)
T5 (320-480ms)
T6 (480-535ms)
Connectivities (C)
(wMNE/PLV)
States (S)
(NMF)
Time (s)
Temporal Similarity
Spatial Similarity
Global Similarity
C1: S1 S2 S3 S4 S5 S6
.81 .10 .09 .20 .43 .01
.82 .80 .58 .41 .45 .58
.81 .45 .34 .30 .44 .29
0.81
C2: S1 S2 S3 S4 S5 S6
.35 .10 .06 .03 .09 .03
.80 .77 .65 .31 .45 .59
.57 .44 .35 .17 .27 .31
0.57
C3: S1 S2 S3 S4 S5 S6
.21 .05 .06 .06 .15 .06
.80 .77 .63 .30 .42 .55
.51 .41 .34 .18 .29 .30
0.51
C4: S1 S2 S3 S4 S5 S6
.06 .85 .07 .24 .15 .12
.79 .76 .62 .31 .44 .53
.43 .80 .35 .27 .29 .33
0.80
C5: S1 S2 S3 S4 S5 S6
.12 .14 .92 .67 .57 .31
.45 .44 .68 .54 .63 .22
.28 .29 .80 .60 .60 .26
0.80
C6: S1 S2 S3 S4 S5 S6
.04 .04 .09 .20 .34 .67
.60 .54 .44 .22 .29 .59
.32 .29 .27 .21 .32 .63
0.63

### Kmeans

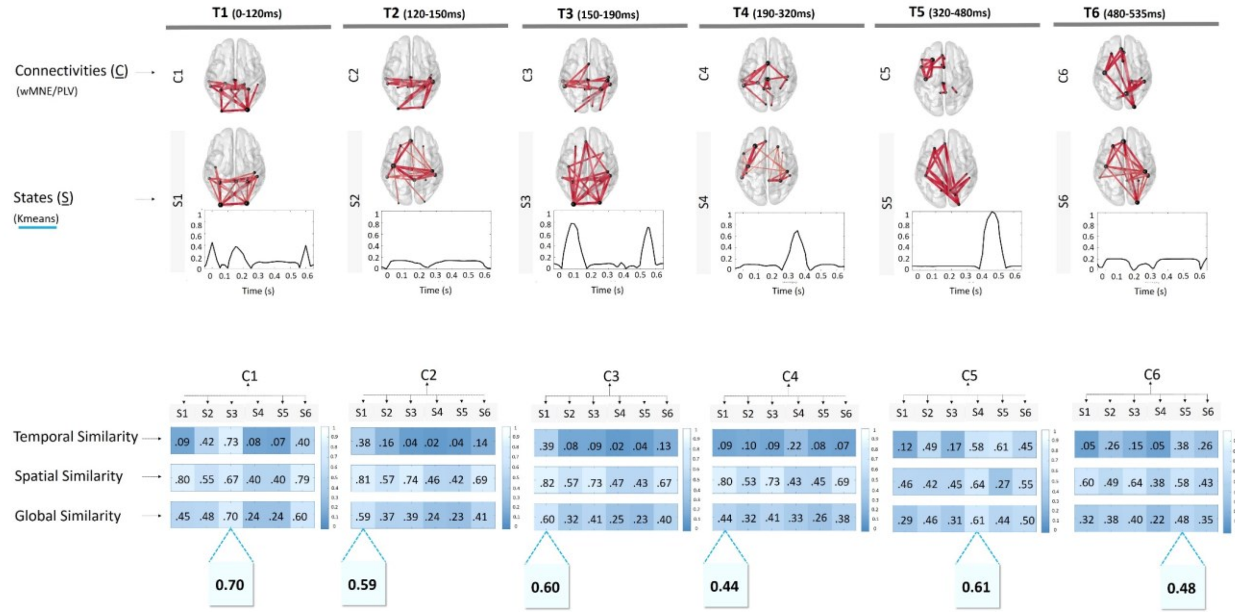


*Figure S3. Detailed results of dimensionality reduction methods (JADE, FastICA, PCA, NMF and Kmeans).*

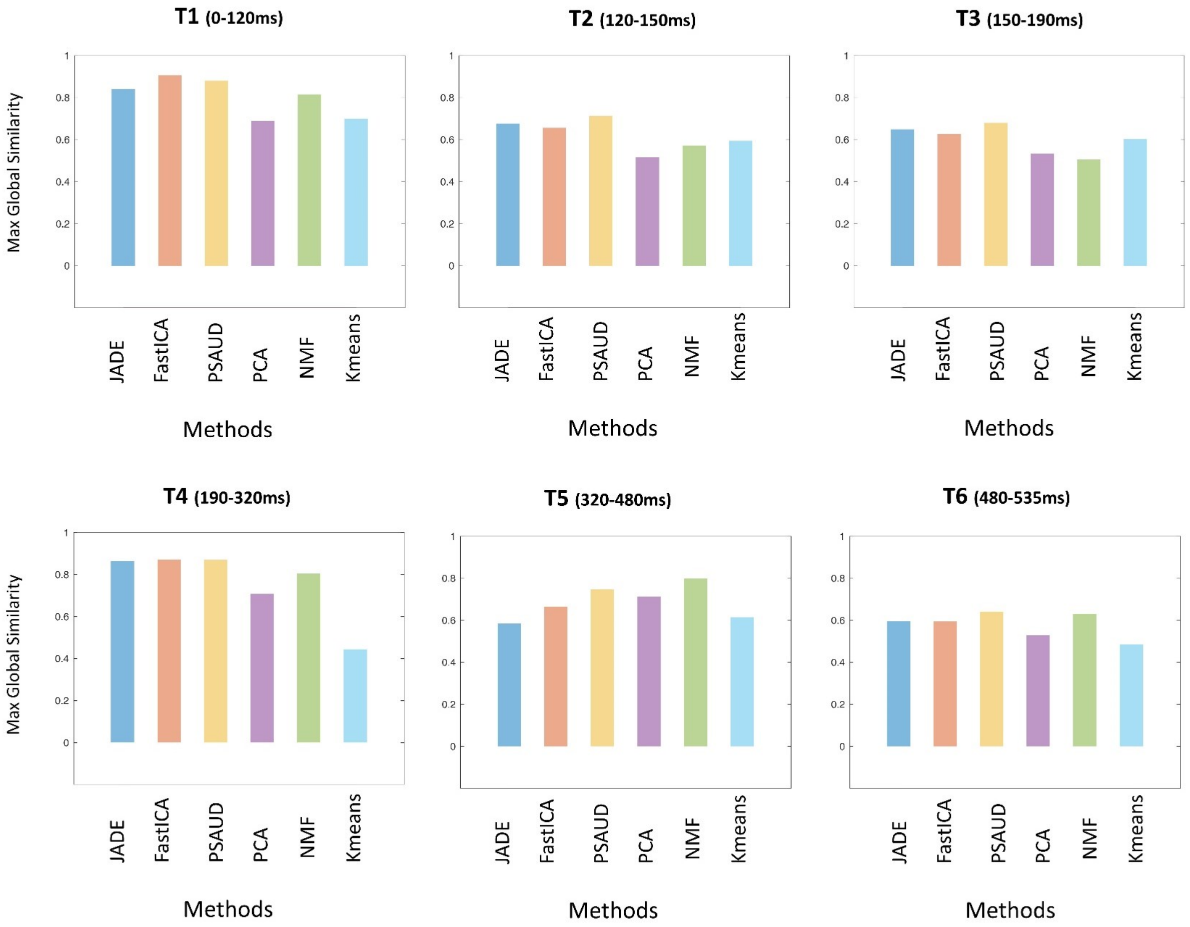


*Figure S4. Maximal Global Similarity results for each dimensionality reduction method at each time intervals. Colors refer to different dimensionality reduction method.*

**A.**

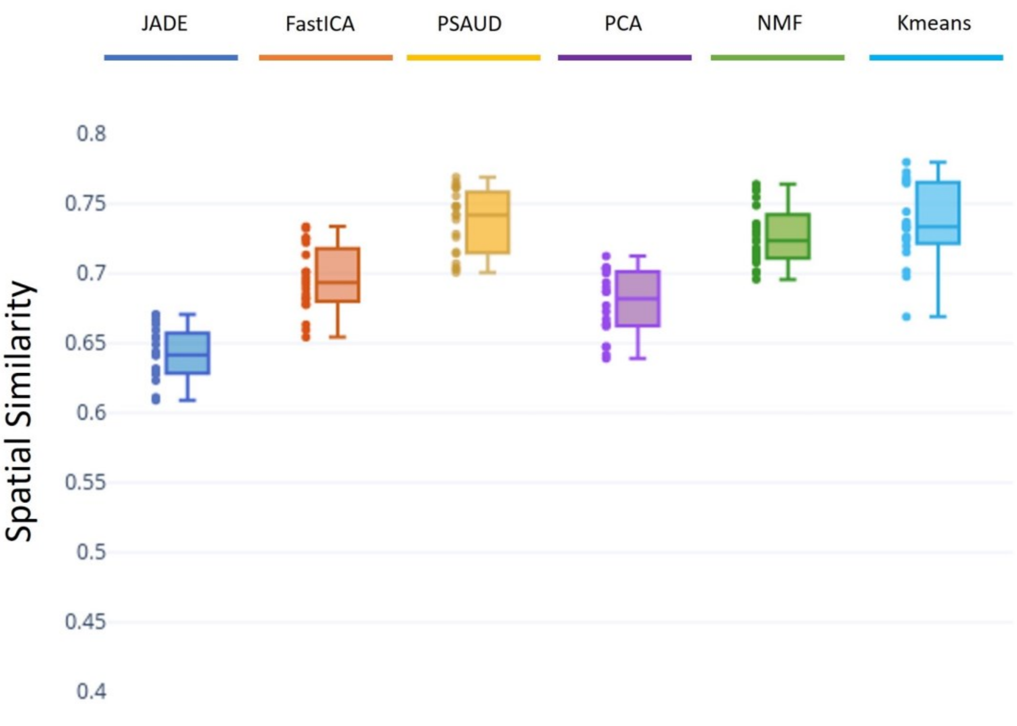


**B.**

| Method 1 | Method 2 | P-value (* for p-value<0.05) |
|---|---|---|
| JADE | FastICA | 6.928 e-10 * |
| JADE | PSAUD | 0 * |
| JADE | PCA | 3.981 e-05 * |
| JADE | NMF | 0 * |
| JADE | Kmeans | 0 * |
| FastICA | PSAUD | 1.793 e-06 * |
| FastICA | PCA | 0.319 |
| FastICA | NMF | 6.233 e-04 * |
| FastICA | Kmeans | 4.212 e-06 * |
| PSAUD | PCA | 1.846 e-11 * |
| PSAUD | NMF | 1 |
| PSAUD | Kmeans | 1 |
| PCA | NMF | 2.036 e-08* |
| PCA | Kmeans | 4.930 e-11 * |
| NMF | Kmeans | 1 |

**C.**

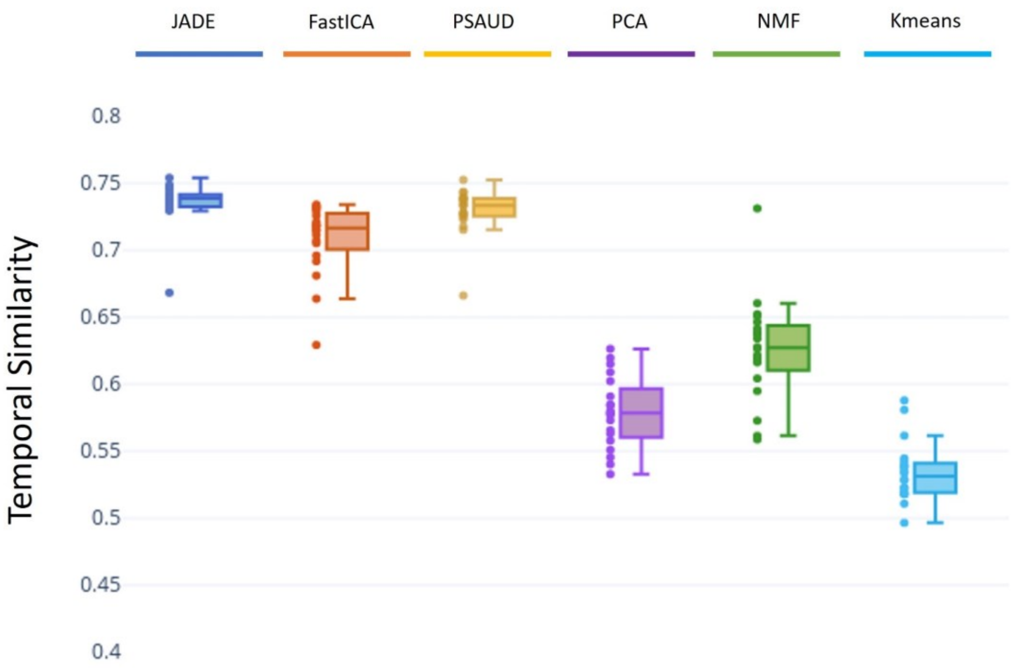


**D.**

| Method 1 | Method 2 | P-value (* for p-value<0.05) |
|---|---|---|
| JADE | FastICA | 0.0195 * |
| JADE | PSAUD | 1 |
| JADE | PCA | 0 * |
| JADE | NMF | 0 * |
| JADE | Kmeans | 0 * |
| FastICA | PSAUD | 0.1848 |
| FastICA | PCA | 0 * |
| FastICA | NMF | 0* |
| FastICA | Kmeans | 0 * |
| PSAUD | PCA | 0 * |
| PSAUD | NMF | 0 * |
| PSAUD | Kmeans | 0 * |
| PCA | NMF | 9.016 e-07 * |
| PCA | Kmeans | 3.843 e-06 * |
| NMF | Kmeans | 0 * |

*Figure S5. Boxplot of spatial similarity (A.) and temporal similarity (C.) distribution over all subjects between each reference state and the matching estimated state for all dimensionality reduction methods. The corresponding p-values of ANOVA statistical test are shown in (B.) and (D.) to evaluate significance differences between methods in both spatial and temporal modes.*

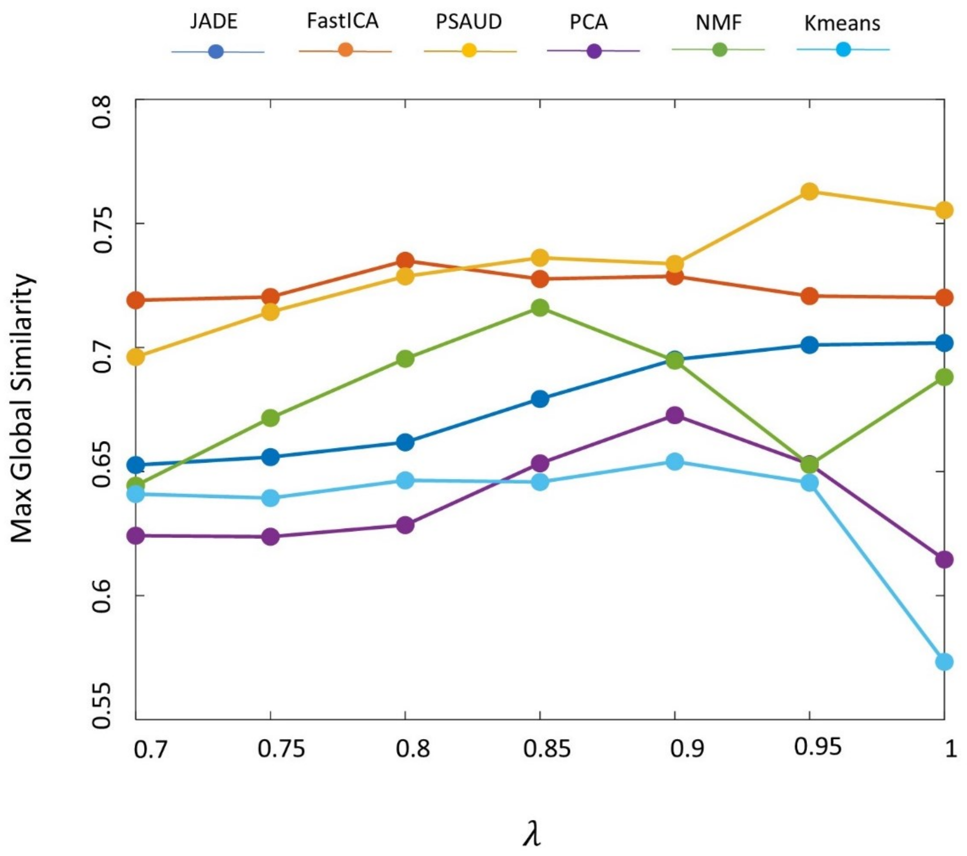


*Figure S6.* Noise variation effect on dimensionality reduction methods performance. The maximal global similarity is computed at the group-level and averaged over all time intervals for each noise level represented by λ value ranged from 0.7 (for the most noisy data) to 1 (for the least noisy data). *Colors refer to different dimensionality reduction method.*
